# Missing the forest for the trees: Do seaweed ecosystems mitigate atmospheric CO_2_ emissions?

**DOI:** 10.1101/2021.09.05.459038

**Authors:** John Barry Gallagher, Victor Shelamoff, Cayne Layton

## Abstract

Global seaweed carbon sequestration estimates are currently taken as the fraction of the net primary production (*NPP*) exported to the deep ocean. However, this perspective does not account for CO_2_ from the consumption of external subsidies. Here we clarify: i) the role of export relative to seaweed net ecosystem production (*NEP*) for a closed system and one more likely open to subsidies; ii) the importance of subsidies by compiling published estimates of *NEP* from seaweed-dominated ecosystems; and iii) discuss their impact on the global seaweed net carbon balance and other sequestration constraints as a mitigation service. Examples of seaweed *NEP* (n = 18) were sparse and variable. Nevertheless, the average *NEP* (−9.2mmol C m^−2^ day^−1^ SE ± 11.6) suggested that seaweed ecosystems are a C source, becoming increasingly heterotrophic as their export is consumed. Critically, mitigation of greenhouse gas emissions was mixed relative to their replacement or baseline states, and where CO_2_ is supplied independently of organic metabolism and atmospheric exchange we caution a sole reliance on *NEP* or *NPP*. This will ensure a more accurate seaweed mitigation assessment, one that does exceed their capacity and is effective within a compliance and carbon trading scheme.

## Introduction

Anthropogenic greenhouse gas emissions (*GHG*) are largely responsible for global warming (Cook et al., 2013). Concerns about warming have led to a call to reduce reliance on the burning of fossils fuels, but also to restore and protect existing natural carbon sinks (UNFCCC, 2015). The most visible of these natural sinks are terrestrial forests, nevertheless, there has been an increasing focus on the advantages and ability of coastal canopy systems to sequester *GHGs*. These are the highly productive saltmarsh, mangroves, and seagrass habitats, which are collectively known as blue carbon ecosystems (Lovelock and Duarte, 2019; McLeod et al., 2011; Nellemann et al., 2009). Unlike terrestrial forests, coastal systems do not readily combust and will continue to sequester carbon down a relatively rapidly accreting sediment column, and at a rate that can respond, and in part determined, by sea-level rise (Lovelock and Reef, 2020). Furthermore, their ability to trap organic imports has resulted in a relatively high carbon sink density, estimated to contribute to half of the total carbon stored in the world’s oceans, despite covering only < 2% of it’s area (Duarte, et al., 2005). Unlike conventional blue carbon systems, seaweeds do not generally support an accreting sediment column. They tend to occur in more exposed rocky reefs and sands where there is little sediment accumulation. As a result, eroded or dislodged seaweed tissue and any excretion of dissolved organic carbon are more typically exported to other coastal and oceanic environments (Duarte and Cebrián, 1996; Krause-Jensen and Duarte, 2016). Nevertheless, seaweed-dominated ecosystems are increasingly being considered an important part of a global carbon mitigation strategy (Bayley et al., 2021; Filbee-Dexter and Wernberg, 2020; Hill et al., 2015; Krause-Jensen and Duarte, 2016; Gallagher, 2014; Krause-Jensen et al., 2018; S. V. Smith, 1981). This is largely because of their considerable global extent, combined with a high rate of net primary production (*NPP*) and potentially high rates of carbon export (~ 43% *NPP*), of which a significant fraction can be sequestered to the deep ocean (~11% *NPP*) (Duarte and Cebrián, 1996; Krause-Jensen and Duarte, 2016).

We contend, however, that the seaweed *NPP* paradigm, which quantifies sequestration as the fraction of seaweed *NPP* exported to the deep ocean, is an incomplete metric of sequestration and by extension mitigation of atmospheric *GHGs*. In this context, sequestration is the rate at which CO_2_ is taken from the atmosphere and stored over climatic scales (e.g. >100 years), whilst mitigation is a measure of the impact on *GHG* emissions should the ecosystem be degraded or lost to an alternative ecosystem state.

The seaweed *NPP* paradigm implicitly ignores the consumption of imported organic subsidies. Indeed, organic subsidies contribute to many wetland systems and some degraded blue carbon ecosystems being rendered net sources of carbon emissions (Duarte and Prairie, 2005). Such imports inevitably result in additional CO_2_ emissions from the stimulation of organic and calcareous metabolism by the seaweed community (Bach et al., 2021; Gattuso et al., 1997). Furthermore, assessing an ecosystem’s mitigation service ultimately requires some estimate of that systems’ carbon balance relative to either a previous baseline state or a future replacement ecosystem. In other words, mitigation services should not be assessed relative to net carbon neutrality but instead, relative to the carbon balance of what would otherwise fill that biological space (Gallagher, 2017; Prairie et al., 2018; Siikamäki et al, 2013; P. Smith, Powlson, Smith, Falloon, and Coleman, 2000). For example, a degraded kelp forest may progress to an alternative state of an urchin barren or turf-dominated assemblage (Edwards et al., 2020; Strain et al., 2014), and *Fucus vesiculosus* assemblages may be replaced by a mussel dominated reef system (Petraitis et al., 2009). Any interventions from a carbon sequestration standpoint will then be dependent on their relative sequestration or emission strengths. However, it is important to also recognise that the choice of an ecosystem carbon sink service must also be constrained by any losses of other ecosystem services such as biodiversity (Villa and Bernal, 2018).

Here we aim to first explore and explain the role of organic subsidies in influencing seaweed ecosystem sequestration relative to the current *NPP* sequestration paradigm. We attempt this by disentangling the components of the ecosystem and its exported net carbon balance: first, for a hypothetical macroalgal system closed to inputs of organic subsidies (Case i), and second, relative to a more usual macro–microalgal ecosystem open to subsidies (Case ii). Second, we assess the importance of subsidies to both the ecosystem’s local and net global carbon balance by compiling published net ecosystem production (*NEP*) estimates before applying export consumption and deposition parameters. These parameters are based on the global *NPP* paradigm model, a compilation average of 30 *NPP* examples across the globe, also previously applied at both continental and (Filbee-Dextor and Wernberg, 2020) and oceanic regional scales (Bayley et al., 2021). Finally, and where available, the expected differences between anthropogenically driven replacements’ local and net global carbon balances are cited to assess where seaweed ecosystem mitigation services lie whilst considering how measurements are constrained by the production of CO_2_ during faunal and floral calcification, and the occurrence of any local upwelling and downwelling processes.

## Materials and Methods

To explore the role of subsidies on seaweed ecosystems, we first partitioned the components of their carbon balance for the ecosystem and its exported material for one system closed and one open to those imported subsidies (Eq. 1 and 3). To gauge the importance of subsidies, published estimates of *NEP* rates were collected from the *Web of Science* database (accessed March 2021) using the following search terms: (macroalga* OR benth*) AND “primary producti*” AND (ecosystem OR community). This search initially identified 2,313 papers, which were subsequently divided and screened by the authors for inclusion based on title and abstract. Only papers that reported, or allowed for, estimates of daily (24 hr) net ecosystem production (*NEP*) of seaweed-dominated communities were used. Results from papers reporting fluxes as oxygen was converted to carbon using a molar photosynthetic and respiratory quotients = 1. It should be noted, that these conversions likely represent a conservatively high estimate as no consideration is given to the potential production of CO_2_ from calcification (Bach et al., 2021; Gattuso et al., 1997). When necessary, the data required to recalculate annual *NEP* from their components were digitally extracted from figures using Graph Grabber™ v2.0.1. We included studies that measured day-time and night-time production/respiration for >1 hr from which we calculated daily estimates using a stated 12:12 day:night ratio (Miller et al., 2009). For one article that reported for 12 hours of daylight only (Miller et al., 2011) we corrected for community respiration rates extrapolated over the night, and at one site, the average *NEP* between various canopy types was weighted using relative biomass (Supplementary material S1, Part 1). Studies that estimated production rates for whole communities based on the summed production rates of individual species were only included if they accounted for both shading by canopy species and respiration of the faunal community (e.g., Miller et al., 2011). Finally, the references contained within the included studies were checked for additional appropriate studies. Data from two papers that measured the *NEP* of *F. vesiculosus* communities from the same sites and depth range in the Baltic Sea and supported almost identical values and were pooled and averaged to minimise any overwhelming influence of this system on the overall mean value across the small pool of included studies (n = 18) (Table 1, Fig. 2). Whilst the pool was relatively small, it resembled a similar sample scale across a range of climatic regions as used in the *NPP* paradigm (n = 30; Krause and Duarte, 2016).

**Table 1.** Net ecosystem productivity (*NEPo*) of seaweed assemblages from the published literature; AEC = aquatic eddy covariance

| Community description | Location | Method | Average $NEPo$ (mmol C m<sup>-2</sup> day<sup>-1</sup>) | Sampling including multiple seasons | Reference |
| --- | --- | --- | --- | --- | --- |
| **Temperate and polar** |  |  |  |  |  |
| Crustose algae/urchins, Fjord | Greenland | AEC | -4.7 | Yes | Attard et al. (2014) |
| *Fucus vesiculosus* | Baltic sea, Finland | AEC | 68.5 | Yes | Attard et al. (2019a) |
| *Fucus vesiculosus* | Baltic sea, Finland | AEC | 66.2 | Yes | Attard et al. (2019b) |
| Brown alga (Phaeophyceae) dominated community | Southern Australia | Chamber | 4.6 | Yes | Cheshire et al. (1996) |
| *Eualaria fistulosa* | Alaska, USA | Chamber | -7.5 | Polar summer | Edwards et al. (2020) |
| Turf algae | California, USA | Chamber | -164.4 | Subtropics winter | Miller et al. (2009) |
| Foliose algae | California, USA | Chamber | -32.64 | Subtropics winter | Miller et al. (2009) |
| *Macrocystis pyrifera* & understory | California, USA | Chamber and growth rates | -8.57 | Yes | Miller et al. (2011) |
| *Macrocystis pyrifera* understory species | California, USA | Chamber | -47.9 | Yes | Miller et al. (2011) |
| *Laminaria pallida* and *Ecklonia maxima* | South Africa | Community carbon balance | -17.7 | Yes | Newell and Field (1983) |
| *Fucus vesiculosus* | Baltic sea, Finland | AEC | 11.2 | Temperate autumn | Rodil et al. (2019) |
| Mixed macroalgae (15-20m) | West Antarctic Peninsula | AEC | -21.8 | Polar summer | Rovelli et al. (2019) |
| *Sargassum horneri* | California, USA | Chamber | -1.8 | Yes | Sullaway and Edwards (2020) |
| **Tropical and subtropical** |  |  |  |  |  |
| *Sargassum* sp. (0.4-3.4m) | Tropical reef, Australia | Open water sampling, other | 10 | Yes | Gruber et al. (2017) |
| *Corallina elongata*, coral reef | Northwest Mediterranean | Open water sampling, other | 20 | Temperate late winter | Bensoussan and Gattuso, (2007) |
| Mixed macrophyte/coral reef mesocosm | Arizona, USA | Open water sampling, other | -3 | Yes | Falter et al. (2001) |
| Mixed macrophyte/coral reef, fringing degraded reef | French Polynesia | Open water sampling, other | 10 | Tropical: winter | Gattuso et al.(1997) |
| Mixed macrophyte/coral reef | Central Red sea | Chamber | 30.4 | Tropical unknown season | Roth et al. (2019) |

### Partitioning the carbon balance

The components of the ecosystems’ local net carbon balance (i.e., *NEP*) are assumed to represent a steady-state over an annual cycle. In this way, seasonality is normalised, although the small number of studies conducted in the growing season likely represent overestimates of annual *NEP,* whilst studies not conducted during the growing season could underestimate annual *NPP*. This is the same level of analysis implicitly used in the global seaweed *NPP* paradigm as a compilation of examples using different methods, at different times, and across different regions (Krause-Jensen and Duarte, 2016). Furthermore, was similarly assumed that any consumption by herbivores, detritivores, or microflora is directed to remineralisation and not an increase in their net biomass or excretion rates. For illustrative purposes, seaweed export is represented as its litter being composed of both the more visible particulate and the less certain fate of its dissolved organic components (Gallagher, 2015; Krause-Jensen and Duarte, 2016). The role of dissolved inorganic carbon is acknowledged but not included because of the uncertainty of it’s production rate and the fate of it’s export (Santos et al., 2021).

#### Case i: The NPP paradigm, a seaweed assemblage closed to imports

This hypothetical system is one where there is no import of organic subsidies for a macroalgal system that dominates primary production. In other words, productivity and respiration contributions from its microalgal assemblage are considered negligible (Fig. 1a). Here, the ecosystems’ net carbon balance (*NEPc*, Eq. 1) is determined between the seaweed assemblage gross primary production (*GPP*) and respiration shared across the alga (*Pr*), and it’s consumption, largely by herbivores (*Hr*) and detritivores (*Dr*) (Duarte and Cebrián, 1996). The remaining production is by inference of a steady state is exported (*E*) to adjacent coastal and oceanic waters. As such, the export term then provides an estimate of the seaweed ecosystems’ potential to sequester carbon and its equivalence with it’s *NEPc* (Eq. 1). However, that sequestration potential will be reduced as it is consumed and remineralised (*Er*) (Krause-Jensen and Duarte, 2016). What remains of the export (*Es*) has been shown to find it’s way to the deep ocean (Fig. 1a), to become a proxy for the seaweeds’ net ecosystem sequestration service (*NES,* Eq. 2). Or expressed more conveniently as the fraction of the seaweeds’ *NPP* that reaches deeper waters (– θ***NPP***, Eq. 2). This then is the implicit rationale behind prominent current estimates (Krause-Jensen and Duarte, 2016).

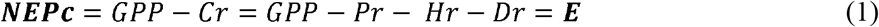

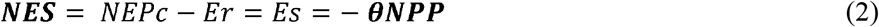

**Fig. 1.**
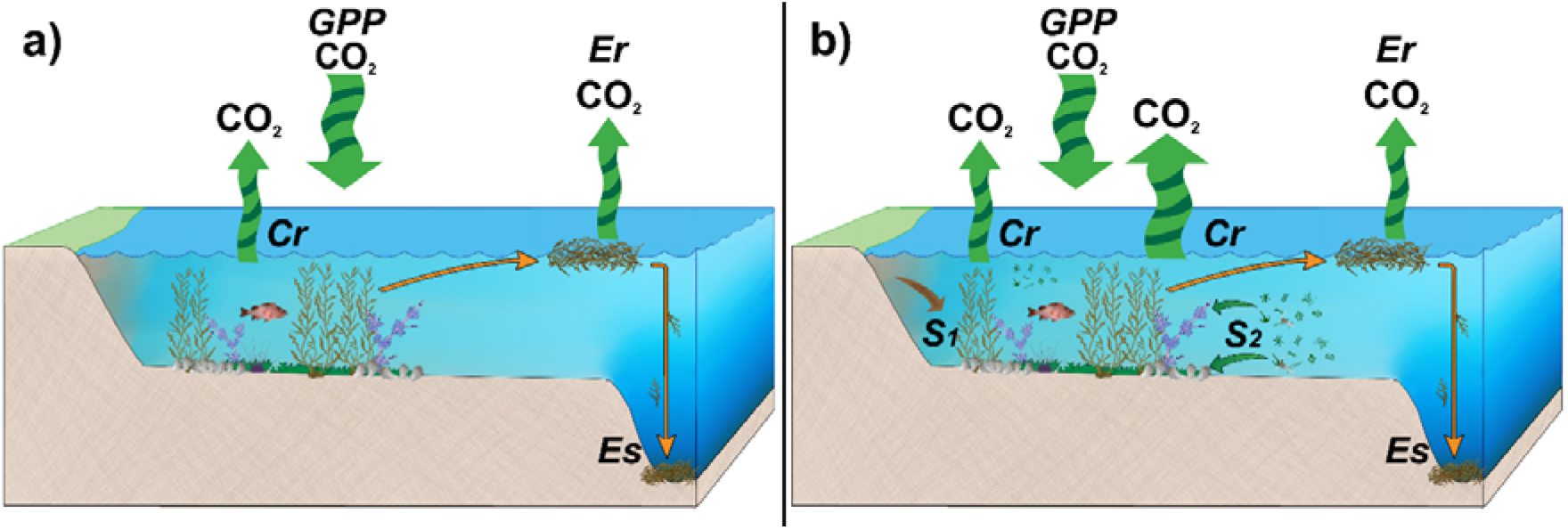
Representation of the components of the net global carbon balance for seaweed ecosystems a) a hypothetical seaweed assemblage closed to the import of organic carbon subsidies, and b) a more representative seaweed–phytoplanktonic ecosystem open to the import of subsidies. Where *Cr* is community respiration partitioned between the algae and the faunal detritivore and herbivore assemblage, *GPP* is gross primary productivity of the primary producer assemblage, *E* is the organic carbon exported from the system, *Er* is the amount of exported organic carbon consumed then remineralised, *Es* is the remaining carbon sequestered in the deep ocean, *S1* represents the supply of any terrigenous organic subsidies and *S2* the organic subsidies supplied from coastal waters, all consumed by the faunal assemblage.

#### Case ii: A seaweed ecosystem open to imports

In reality, though, these seaweed-dominated systems are open to organic imports (Foley and Koch, 2010; Robert J. Miller and Page, 2012; Zuercher and Galloway, 2019) and support an interactive phytoplanktonic assemblage that shares the ecosystems’ *NPP* (Borum and Sand-Jensen, 1996; Miller et al., 2011), and (Fig. 1b). As these imports are consumed by the faunal community, they subsidise the release of CO_2_ (*Sr*) by further stimulating their organic and calcareous metabolism, and thereby lowering the net ecosystem production (*NEPo*) (Eq. 3). As for Case i, *Er* can be approximated as the amount of carbon exported from the seaweed assemblage. The relative contribution from phytoplankton export does not appear to be significant (see Supplementary material, Part 2).

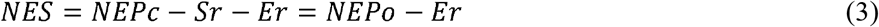

It can be appreciated from Eq. 3 that the additional respiration from the consumption of subsidies (*Sr*) is likely to be substantial when annual *NEPo* is either in balance or indeed, heterotrophic (i.e. *–NEPo*). Moreover, the greater the influence of *Sr* on the ecosystems’ carbon balance, the greater the over-estimation of an *NES* based solely on the fraction of the seaweeds *NPP* (*–* θ*NPP*) exported to the deep ocean (Case i).

## Results

Studies with year-round sampling often showed strong seasonal effects with lower *NEP* values in the cooler/shorter daylength seasons and higher *NEP* values in the warmer/longer daylength seasons (i.e., increased growth) (Attard et al., 2014; Attard et al., 2019a; Karl M. Attard et al., 2019b; Cheshire et al., 1996; Falter, et al., 2001; Miller et al., 2011; Sullaway and Edwards, 2020). Additionally, over annual cycles, *NEP* rates within the warmer and tropical–subtropical ecosystems supporting also higher light intensities, appeared to be greater than their more heterotrophic temperate–polar counterparts (*p*(same median) < 0.04 Mann-Whitney U) (Fig. 2). Overall, the examples, largely including average annual estimates, varied substantially around a heterotrophic mean (Fig. 2) of −9.2mmol C m^−2^ day^−1^ (SE ± 11.6). The stand-out exceptions were *F. vesiculosus* supporting highly autotrophic annual *NEP* rates and extreme heterotrophy demonstrated by a turf dominated assemblage (Miller et al., 2009) (Fig. 2, Table 1).

**Fig. 2.**
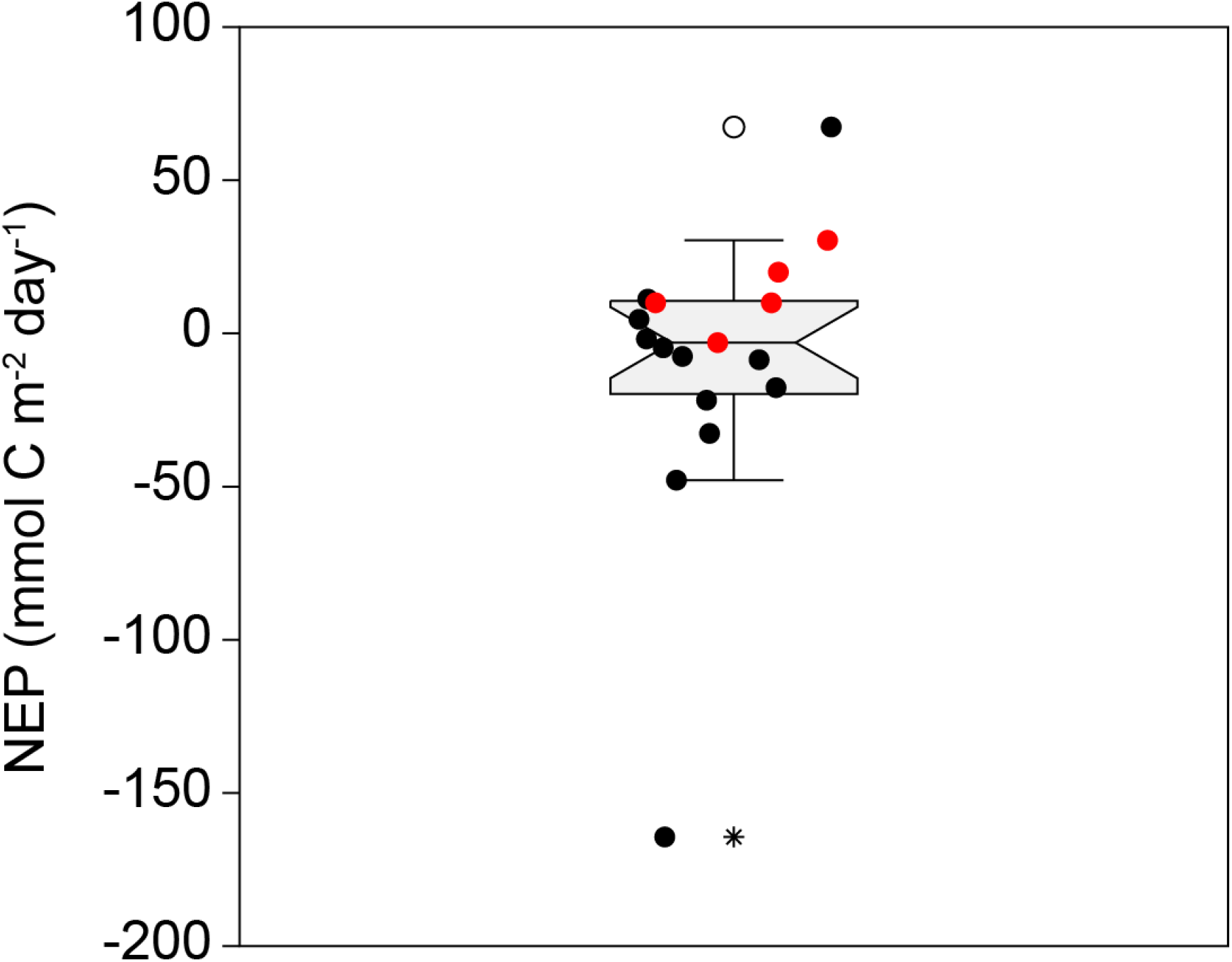
Box and whisker plot for seaweed ecosystem net ecosystem production (*NEPo*) extracted from the literature for polar to tropical communities (• temperate to polar systems; • subtropical to tropical systems). Outliers are represented by the symbols (⁕) and (O). The top outlier is the average across two *F. vesiculosus* studies from in the same area and depths of the Baltic Sea, with the central static representing the median of the distribution.

### Seaweed ecosystems’ global carbon balance

Along with net ecosystem production, an account must also be made of the amount exported seaweed production that is consumed and subsequently remineralised during export to the deep ocean. This is implicit in the *NPP* paradigm calculation as the fraction of *NPP* exported that remains after consumption. It can be taken as the difference between estimates of the average fraction of *NPP* exported (43%) and the remains to the deep ocean (11%) as previously used across global, continental, and oceanic island scales (Bayley et al., 2021; Filbee-Dextor, 2020; Krause and Duarte, 2016) (Eq. 4).

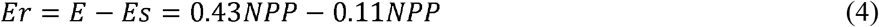

By substituting Eq. 4 into Eq. 3, the *NES* for seaweed ecosystems open to the import of subsidies then becomes the difference between the measured *NEPo* and 32% of the seaweeds *NPP* (Eq. 5).

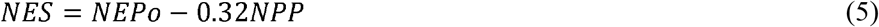

We can then substitute our mean *NEPo* value (−9.2 mmol C m^−2^ day^−1^) and the mean *NPP* for seaweed systems around the globe (n =30; Krause-Jensen and Duarte, 2016), to estimate the net ecosystem service (*NES*) (Eq. 6).

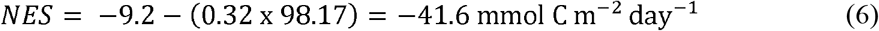

To clarify, such a calculation is only intended to illustrate pertinent concepts and relies on whether the *NPP* sample mean (n = 30) lies close or within the same part of the population distribution as our *NEPo* trimmed compilation (n = 17). Like the *NPP* paradigm, it does not necessarily provide an accurate estimate of global seaweed sequestration. There are also regional bathymetric, climatic time scales, and species effects to be addressed as the science progresses. Nevertheless, like the *NPP* paradigm, the extent of the net global balance identifies an important global carbon vector. In this case, the large differences between the two conceptual models provide a more considered insight on the likely extent of additional consumption of exported material on the seaweeds’ global net carbon balance, be it large or small relative to the global sample average *NEPo*.

## Discussion

### Seaweed carbon balances

It appears that on the whole, seaweed ecosystems are substantially impacted by the consumption of organic subsidies to the extent that on average they appear to be heterotrophic (−9.2 mmol C m^−2^ day^−1^) at local scales. Furthermore, their average global carbon balance becomes increasingly a carbon source to the water column by accounting for remineralisation of their exported production (*cal* −40.6 mmol C m^−2^ day^−1^, Eq. 6), and not a global sink (*cal.* +10.80 mmol C m^−2^ day^−1^, Krause-Jensen and Duarte 2016). Indeed, our estimate suggests that the average seaweed *NEPo* would need to exceed 31.41 mmol C m^−2^ day^−1^ (i.e. the sum of 0.32 and 98.17, Eq. 6) just to maintain a global carbon balance. For many seaweeds supporting a *NEPo* sufficient to overcome the amount of export remineralised appear not to be likely (Table 1). However, this does not exclude other seaweeds such as the temperate *F. vesiculosus* and tropical–subtropical examples that appear to support more autotrophic regimes even during the winter period (Fig. 2, Table 1). Whether this is because of a relatively large *NPP* or smaller import and consumption of subsidies is not clear.

The current research on seaweeds remains mostly restricted to coastal benthic systems. Nevertheless, the *NPP* paradigm has also applies to natural floating species (Bach et al., 2021) and floating rope aquaculture (Duarte et al., 2017; Chung et al., 2011). As for benethic systems as far as we are aware the paradign has not been tested, with appropriate data restricted to a single study from the Yellow Sea, China (Jiang et al., 2013). The study calculated the atmospheric CO_2_ flux from annual changes in *p*CO_2_ within the water columns (34.85 ± 17.46 mmol C m^−2^ day^−1^). While the value was significantly greater than sites adjacent to the seaweed arrays (24.17 ± 14.14 mmol C m^−2^ day^−1^) the difference was reduced towards carbon neutrality when compared to the reported baseline values for the area (32.71 ± 17.23 mmol C m^−2^ day^−1^). Nevertheless, their overall mitigation services are likely to be significant given that export as in harvesting, is conceivably greater than for natural systems (Chung et al., 2011). Information on natural floating *Sargassum spp.* ecosystems; however, is confined to *Cr* and *GPP* components determined from two separate studies of *Sargassum natans* from the oceanic and neritic waters south of Bermuda and the NW Atlantic shelf respectively (Lapointe, 1995; Smith et al., 1973). Together, the *Cr* for neritic and oceanic regions appeared to be more than 3 to 5 times larger than their *GPP* respectively (Supplementary material S1, Part 3), suggesting that subsidies also play a major role in constraining the *NEPo* of those ecosystems.

### Mitigation services

Critically, estimates of a seaweed systems’ global carbon balance (Eq. 6) in isolation, while valuable, requires comparisons of global balances of their actual or potential alternative replacement states (Gallagher, 2017; Siikamäki et al., 2013; Smith et al., 2000). However, *NEPo* measurements for kelp replacements such as barrens and turfs-dominated systems are limited, and the results mixed. Turf ecosystems can support a *NEPo* carbon balance of around −164.4 mmol C m^−2^ day^−1^ (Table 1 (Miller et al., 2009)). This is significantly more heterotrophic than the previous mixed *M. pyrifera* assemblage (−8.57 mmol C m^−2^ day^−1^) from the same region (Table 1 (Miller et al., 2011)). In contrast, urchin barrens across many sites within a polar region appear to be moderately heterotrophic. On average their *NEPo* range from −4.76 mmol C m^−2^ day^−1^ (SE ± 1.35) (Attard et al., 2014) to −3.75 mmol C m^−2^ day^−1^ ^(^SE ± 10.56) (Edwards et al., 2020). These are only marginally less heterotrophic than the kelp forest counterparts (−7.5 mmol C m^−2^ day^−1^ SE ± 7.7) from similar environments (Edwards et al., 2020). For the highly autotrophic *F. vesiculosus* ecosystems, their potential to sequester carbon (66.2 and 68.4 mmol C m^−2^ day^−1^ Table 1) may be amplified when considering the *NEPo* carbon balance of the mussel reef replacement in the same area (−39.5 mmol C m^−2^ day^−1^) (Attard et al., 2019a). Furthermore, further differences in their global net carbon balances (i.e. *NES,* Eq. 5) are unlikely to be great. The net primary production within and between coastal seaweed and phytoplanktonic ecosystems appear to converge (Borum and Sand-Jensen, 1996) along with the amount of the export remineralised (Supplementary material S1, Part 2).

### Other limitations: Inorganic carbon supply and outwelling

We have primarily focussed on the organic carbon balance over a more recent consideration of dissolved inorganic carbon exported as a long-term dissolved sequestration pool, described as outwelling (Santos et al., 2021). There is, however, an aspect of this outwelling that has not yet been addressed. This is the impact of an acidifying ocean and turbulence between a vegetated and non-vegetated system on the dissolution of their edaphic calcareous sands and fauna. Turbulence together with ocean acidification can significantly increase the dissolution of calcium carbonate and conceivably increase the amount of bicarbonate outwelling (Eyre et al., 2014). In contrast, photosynthesis within a seaweed canopy can significantly reduce acidification and turbulence (Morris et al., 2019; Murie and Bourdeau, 2020). In other words, the canopy is reducing the outwelling sequestration pool relative to a non-vegetated alternative or baseline system. We now have a possible situation where the non-vegetated system is the preferred carbon sequestration sink. However, maintaining or transiting to such a system may not be justified if it is also accompanied by smaller biodiversity (Villa and Bernal, 2018) or arguably, the loss of other natural capital services.

The *NEPo* carbon balance is a measure of CO_2_ flux to or from the atmosphere for enclosed and semi-enclosed systems (Prairie et al., 2018). In open coastal waters, however, CO_2_ can be supplied independently of atmospheric exchange and organic metabolism. Most notably, from geostrophic currents and upwelling (Ikawa and Oechel, 2015; Thorhaug et al., 2020), as well as faunal (Gattuso et al., 1997) and algal calcification notably the extensive production from seaweed *Halimeda spp* (Borowitzka and Larkum, 1976). These additional sources can conceivably not only not affect atmospheric exchange independent of the *NEPo*, but also invalidate *NEP* and *NPP* concepts as processes driven by CO_2_ sequestered from the atmosphere. Under such conditions, assessments will likely require additional resources to measure atmospheric exchange between the seaweed ecosystem and its replacement, from the same area. A combined understanding of *NEPo* and the fate of local export appear to be the prerequisites necessary for a predictive capacity to fully assess a seaweed ecosystems’ capacity to mitigate *GHG* emissions.

## Future research and conclusions

Seaweed ecosystems may not be the significant sequesters of global carbon that they were previously thought. There are several data gaps and conceptual shortcomings that still need to be addressed, including 1) additional measurements of seaweed *NEPo* over annual cycles; 2) and comparison of these measurements relative to the local alternative or degraded state; 3) further understanding of organic subsidy supply and consumption; 4) estimates of atmospheric flux of CO_2_ to disentangle any physical from the biological divers of atmospheric exchange; 5) measurements of exported production and sequestration at local scales. Until then, robust assertions of carbon sequestration and mitigation by seaweeds appear premature and should be interpreted with prudence. It must also be noted that such overestimates when presented as important at global scales are not always benign. This is particularly the case when considering a carbon credit offset and trade scheme (Johannessen and Macdonald, 2016; Repetto, 2013). Carbon credits may become more expensive for polluters to compensate their emission above their cap and increase GHG emissions above the sequestration capacity of the ecosystem. Finally, and most importantly, irrespective of the role that seaweed-dominated ecosystems play in carbon mitigation of *GHGs*, they should remain highly valued for the vast array of critical ecosystem services they provide, including their incontrovertible support of coastal productivity and biodiversity.

## Acknowledgments

We are grateful to Dr John Gibson for his comments on the carbon balance construct and to Mr Chuan Chee Hoe for his assistance with the illustrations

## Declarations

### Availability of data and material

All data is contained found with the text of Table 1 with details of any recalculations to obtain the data and additional supporting analysis in Supplementary Information (S1)

### Authors’ contributions

Authors contributed to the study conception [John Barry Gallagher] and [Victor Shelamof] with contributions to insights on alternative states [Cayne Layton]. Authors contributed to the collection and processing of the data [John Barry Gallagher], [Victor Shelamof] and [Cayne Layton], and data synthesis [John Barry Gallagher]. The first draft of the manuscript was written by [John Barry Gallagher] and all authors commented on previous versions of the manuscript. All authors read and approved the final manuscript

