## Supplementary Materials for "Missing the forest for the trees: Do seaweed ecosystems mitigate atmospheric CO_2_ emissions?" (from another version of this preprint: version 1, 2021-09-06)

### Missing the forest for the trees: A reappraisal of global seaweed carbon sequestration

### Introduction

The following additional information about the main article is in three parts:

- Part 1 sets out the details of the net ecosystem production (NEP) recalculated from the ecosystem’s carbon balance components reported by Miller et al. (Miller, Reed, & Brzezinski, 2011) for the monitoring site MC supporting both the understory but dominating *Macrocystis pyrifera* assemblage, and near adjacent to site MR supporting the canopy understory assemblage without *M.* *pyrifera.*
- Part 2 provides more details behind the assumption that the fraction of a seaweed ecosystem’s export is largely the macrophyte and not its phytoplankton assemblage
- Part 3 sets out the calculation and data used to calculate NEP for the *Sargassum sp* occupying parts of the Sargasso sea. This was taken from two different studies of the same species and sites that separately measured gross primary production (GPP) and community respiration(Cr) (i.e., NEP = GPP – Cr)

### Part 1

Miller et al. (Miller et al., 2011), reported benthic chamber measurements for the understory macroalgae net primary production (NPP) community respiration (Cr), that is, the respiration for faunal and macroalgal assemblage, for both MR and MC sites across seasons as expected for 12 hrs of daylight.,In addition to benthic chamber understory measurements the large canopy *M*. *pyrifera* NPP was measured for the MD sites using from monthly average aerial growth rates of organic carbon biomass (i.e. the result night and day growth).

#### MR site

To recalculate the annual seasonal average NEP for site MR (labelled as NCP by Miller et al. 2011) for a complete day and night period, it was assumed that the daytime community respiration rates remained the same during the night

Data

All data were extracted digitally from Figures using Graph Grabber 2.0.1.

Annual average understory NEP = 20.67 mmols C m^-2^ day^-1^ over 12 hrs daylight

Average seasonal understory Cr = -116.48 mmols C m^-2^ day^-1^ (minus denotes a CO_2_ source)

*Calculation*

Annual average understory NEP = NEP_12hr_ – Cr_12hrs_

**NEP = (20.67/2) - (-116.48/2) = -47.91mmols C m^-2^ day^-1^**

Where the annual average net CO_2_ emitted over 12 hrs is NEP_12hr_, and the additional amount of CO2 emitted by community respiration over the evening is Cr_12hrs_.

#### MC site

### Data

All data were extracted digitally from Figures using Graph Grabber 2.0.1.

Annual average understory NEP = -66.48 mmols C m^-2^ day^-1^ over 12 hrs daylight

Average seasonal understory Cr = - 66.48 mmols C m^-2^ day^-1^ (minus denotes a CO_2_ source)

Average annual *M. pyrifera* NPP = 44.2 mmols C m^-2^ day^-1^

Understory macroalgal respiration as a fraction of their NPP = 0.37 (Gerard, 1986).

Fraction of annual average understory to *M. pyrifera* biomass within the MC site = 0.048 and 0.952 respectively.

To recalculate the annual seasonal average NEP for understory and *M*. *pyrifera* assemblage the calculation was in three parts:

***First****,* the understory NEP region was calculated in the manner for the MR site (see above).

**NEP_understor_ = (-55.83/2) - (-66.48/2) = -61.16 mmols C m^-2^ day^-1^**

***Second***, the annual average NEP for the *M*. *pyrifera region* was calculated using the difference between the *M*. *pyrifera* daily NPP annual average and its daily faunal community respiration from its associated detritivores (Dr) and herbivores (Hr), that is:

NEP_Mpyr_= NPP – (Dr + Hr).

The estimates for faunal respiration were calculated by subtracting the *M*. *pyrifera* respiration (Pr) as that fraction of its NPP (0.37) from data within Gerard (Gerard, 1986) for the same sampling region and depths.

Cr = Pr + Dr + Hr

Faunal respiration Dr + Hr = Cr - 0.37NPP = 66.48- (0.37 x 44.2) = 50.16 mmols C m^-2^ day^-1^

From which:

**Annual average NEP_Mpyr_ = 44.2-101.3= -5.93 mmols C m^-2^**

Again, this assumes that the community Cr measured during the period of 12 hr of light was the same over the night.

***Third,*** the annual NEP for the understory and M. pyrifera regions of the site we combined as their weighted average based on their relative biomass over an annual cycle

**Annual NEP_MC_ = (-5.93 x 0.952) = (-61.16 x 0.048) = -8.5**

### Part 2

The calculation for global average sequestration rates for macroalgae as set out in the article, also assumes that the fraction of the ecosystem’s export is largely the macrophyte and not its phytoplankton assemblage (Fig. 1b main text). We believe at this level of analysis this is a reasonable assumption for open coastal systems. These areas not impacted by nutrient eutrophication where the phytoplankton assemblage may dominate the ecosystem *NPP* over the macrophyte (Borum & Sand-Jensen, 1996). For example the lush *Macrocystis sp*–turf ecosystems off the Californian coastline, the phytoplankton *NPP* to the total autotrophic assemblage can be calculated to average around 0.21 (Miller et al., 2011). By taking lessons from the fate of autotrophic production of coastal phytoplankton (Duarte & Cebrián, 1996), the difference between the amount exported (0.16 NPP) and stored in sediments (3.9 NPP). Then the average amount of phytoplankton export respired is around 12.4% of their NPP, an additional contribution of less than 0.025 to the exported respiration rate in Eq. 5 of the main text (i.e., 0.32 + 0.025NPP).

### Part 3: Free-floating seaweeds

Along with benthic coastal seaweed ecosystems, a similar argument based on the fraction *NPP* exported has been made for free-floating seaweeds ecosystems for the extensive Great Atlantic Sargassum Belt (GASB) (Bach et al., 2021). The *Sargassum sp* associated with the GASB also supports a vibrant animal and epibiont community, which has also to be open to planktonic subsidies. What fraction of organic subsidies support its animal and epibiont community is not clear? As far as we are aware there have been no measurements of *NEP* on Sargassum sp, or any natural free macroalgae*.* Nevertheless, the separate components of *NEP*, that is, community respiration Cr and the gross primary production (GPP) have been measured in two separate studies. The studies were based on the same species (*Sargassum natans*) and located within the same oceanic and neritic waters south of Bermuda and the NW Atlantic shelf respectively, but around 10 years apart (Lapointe, 1995; Smith, Burns, & Carpenter, 1973). Neritic and oceanic community respiration (*cal^[1]^* 7.28 ± 0.48 and 5.25 ± 0.16mg C gdrywt^-1^ hr^-1^) respectively was more than 3 to 5 times larger than their typical *GPP* respectively (2.38 ± 0.52 and 1.17 ± 0.65mg C gdrywt^-1^ hr^-1^ respectively). Such a difference suggests that these systems are likely to be heterotrophic, or at least no more than close to a carbon balance when considering the phytoplankton production shaded by the surface canopy (Bach et al., 2021).
